# Risks in signal processing pipelines influencing the estimation of phase dependency for EEG-TMS

**DOI:** 10.1101/477166

**Authors:** Robert Guggenberger, Maximilian Scherer, Alireza Gharabaghi

## Abstract

Phase-dependency of cortico-spinal excitability can be researched using TMS-EEG. Due to the large artifact, non-causal filters can smear the TMS artifact and distort the phase. However, causal filters can become biased by too high filter orders or uneven pass-bands. We explored the influence of different signal processing pipelines on the estimation of the optimal phase. This exploration involved performing two simulation studies. In the first, we simulated two different phase-dependencies (uni-versus bimodal) and sought to recover them with two distinct approaches that have previously been described. In the second, we specifically explored how filter parameters (e.g., order, pass-band) biased the phase estimation. On the basis of these findings, we propose using up-to-date toolboxes, re-running scripts after software updates and performing simulation studies in parallel to safeguard the analysis pipeline of empirical studies.

## Introduction

The excitability of neuronal populations depends on their current state (Izhikevich, 2007). The use of non-invasive measurements to monitor the current state of neuronal populations is an important step towards implementing brain-state-dependent transcranial stimulation for research and therapy. This is feasible with power-based approaches. For example, desynchronization of sensorimotor power, as picked up by electroencephalography (EEG), increases cortico-spinal excitability (CSE), as measured by the motor-evoked potential (MEP) in response to transcranial magnetic stimulation (TMS; Takahashi et al., 2018; Takemi et al., 2013; Kraus et al., 2016, 2018). Similarly, transcranial alternating current stimulation (tACS), which is assumed to synchronize neuronal populations (Zaehle et al., 2010), is able to increase CSE. Such knowledge has already been used to implement interventions for clinical application (Gharabaghi et al., 2014).

Most TMS devices that apply short-lasting single pulses can be triggered with very short latencies. Such high temporal precision makes it possible to probe the state-dependency of CSE not only with respect to the rather slow-changing measure of oscillatory power but also with regard to the relatively fast-changing phase of an oscillation. Research has suggested that the use of TMS at different phases of sinusoidal signals applied via transcranial alternating current stimulation (tACS) discloses such phase-dependency of CSE (Raco et al., 2016; Guerra et al., 2016; Nakazono et al., 2016; Fehér et al., 2017; Schilberg et al., 2018).

The step from exogenous entrainment of oscillations via tACS to reading the current endogenous phase of an oscillation from EEG or EMG recordings entails several challenges. Unlike a tACS signal, the signal-to-noise ratio of non-invasive electrophysiological recordings is low. The amplitude and phase of oscillations are non-stationary and might be sinusoidal only in approximation (Cole and Voytek, 2017). In an attempt to address this non-stationarity, the construction of dedicated hardware might become necessary for fast processing. However, despite using dedicated hardware, and even if performed only when the oscillation amplitudes are high, real-time estimation of an oscillations phase might have a standard deviation of about 50° (Zrenner et al., 2017).

Post-hoc analysis of TMS, applied initially at random and subsequently probed for possible phase-dependencies, therefore remains an important method of research (van Elswijk et al., 2010; Keil et al., 2013; Khademi et al., 2018). Nonetheless, this approach still requires the stringent design of a signal processing pipeline with a special focus on the filter design. For example, non-causal filters may smear the TMS artifact and distort the phase of an oscillation prior to the TMS pulse. Furthermore, when transferred to real-time applications, only causal filters are feasible. In this study, we show how methodological differences in the design of causal filters affect the estimation of the phase dependency. This is not simply a methodological question. For example, with regard to the phase-dependency in the oscillatory beta-band, it is unclear as to whether there is only one maximum of CSE, found in the rising phase of an oscillation (Khademi et al., 2018; van Elswijk et al., 2010), or whether there are two maxima of CSE, i.e., at the peak and trough of an oscillation (Keil et al., 2013). Using simulated data, we show how two different approaches of phase estimation can result in such conflicting findings, even when based on identical data. Subsequently, we explore the general influence of the filter order and bandwidth on recovering the phase of maximal CSE. Finally, we discuss policies for minimizing risks in the design and implementation of signal processing pipelines for estimation of phase dependency.

## Material and Method

All simulations were performed and visualized with Anaconda Python 3.6.5 on Linux Mint 18.2, employing SciPy 1.1.0, NumPy 1.14.3, Seaborn 0.8.1 and Matplotlib 2.2.2. The script to create simulated dependencies is available online (https://osf.io/mgu4h/).

### Simulation 1

We simulated data with two dependencies between oscillatory phase and MEP amplitude. One was bimodal, i.e., with two maxima of MEP at peak and trough of the oscillation. The other was unimodal, i.e., with only a single maximum at the rising flank. Both dependencies were sampled, filtered and processed in accordance with the methods reported previously (Keil et al., 2013; van Elswijk et al., 2010).

#### Data model

In the dependency models, data was simulated as a 18 Hz cosine signal, and subsequent analyses were performed in accordance with two previous reports. The first approach involved estimating the phase in analogy to Keil and colleagues (Keil et al., 2013) on the basis of a Hilbert transformation. We simulated the data with a 2 kHz sampling rate and duration of ±1.5 s, and included three cycles, terminating 5 ms before the TMS pulse. The second approach entailed estimating the phase in analogy to Elswijk and colleagues (van Elswijk et al., 2010) on the basis of a discrete Fourier transformation. We simulated the data with a 10 kHz sampling rate and duration of ± 1.1 s, and included two cycles prior to the TMS pulse.

#### MEP dependency

We simulated two different dependencies between the oscillatory phase and MEP amplitude: maximum MEP amplitude (i) with a unimodal pattern, i.e., at the *rising* phase of 18 Hz oscillations, (ii) with a bimodal pattern, i.e., at the peak and trough of 18 Hz oscillations. We simulated a log-linear relationship with base 50 to account for the log-normal distribution of MEPs. The two dependency models are presented in figure 1A.

**Figure 1.**
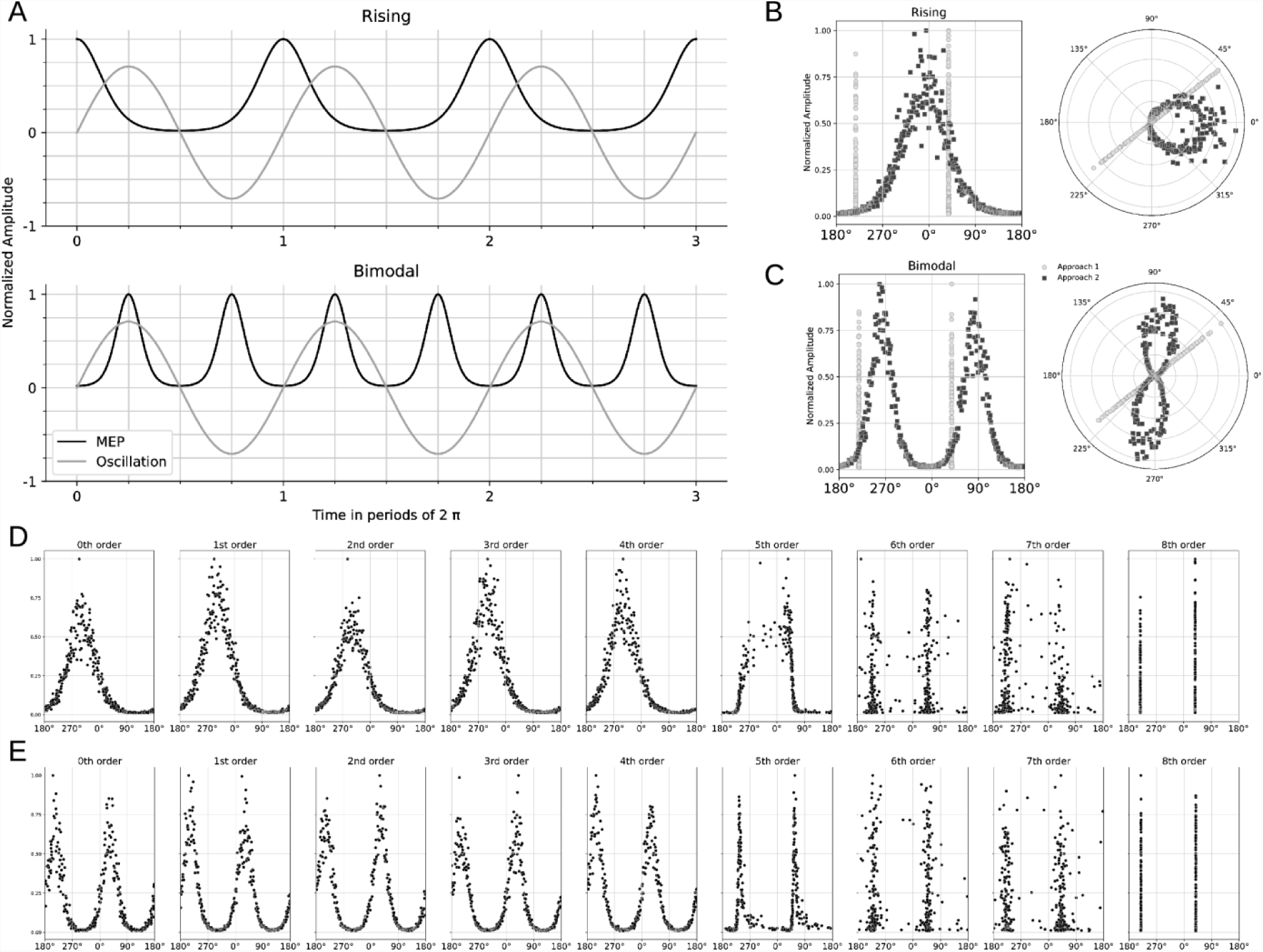
Recovery of uni- and bimodal models depending on the approach applied. ***A***, Upper row shows unimodal phase-dependency with maximum MEP amplitude (black line) at the rising flank of oscillatory activity (gray line). Lower row shows bimodal phase-dependency with maximum MEP amplitude (black line) at the peak and trough of oscillatory activity (gray line). ***B*** Unimodal phase-dependency which is not recovered by narrow-band filtering followed by Hilbert transformation (approach 1, light dots) but by broad-band filtering followed by discrete Fourier transformation (approach 2, dark dots).***C*** Bimodal phase-dependency which is recovered by approach 2 (dark dots) and in a distorted way by approach 1 (light dots). Left columns are scatter plots with phase on the x-axis and recovered MEP amplitude on the y-axis. Right columns show the same data as a polar plot on the unit circle. **D/E** The recovery depending on filter order for approach 1, i.e., narrow-band (17-19 Hz) filtering followed by Hilbert transformation with different filter orders (from zero to 8^th^) for the unimodal (E) or the bimodal (F) model. Lower filter orders can distinguish between bimodal and unimodal dependencies of MEP amplitudes. Note that, due to the estimation of the phase 5 ms prior to the TMS-pulse, a phase-shift occurs even without filtering.

#### Filter Order

Simulated data epochs were bandpass filtered forward in time with a Butterworth filter with orders from 0 to 8, and for two different frequency bands depending on the approach, i.e., 10-400 Hz (van Elswijk et al., 2010) or 17-19 Hz (Keil et al., 2013).

#### Assessment

We visualized the simulation results as polar-linear and linear-linear scatter plots between oscillatory phase and MEP amplitude. To achieve a well-resolved and uniform sampling across the unit circle, the relationship for each integer phase was sampled from 0° to 360°. For the simulation, we added a small white noise term (with 0.05 standard deviation) to prevent points from overlaying and to reduce any possible bias due to numerical errors.

### Simulation 2

We simulated data as above, but with one dependency mode only, i.e., peak of CSE would be at the rising flank of an oscillation at 18 Hz. Signals were simulated with a sampling rate of 1000 Hz, for the duration of ± 1.5 s around the TMS for filtering, and included three cycles for subsequent phase estimation. The signal was simulated without noise.

#### Data processing

We filtered the data by exploring the influence of four parameters, i.e., *estimation method, filter order, filter bandwidth*, and *filter center*. The phase was estimated using either Hilbert or Fourier transformation as described above. Additionally, we systematically increased filter order up to an order of 8, and decreased the filter bandwidth. This was achieved in two ways, with the bandwidth (i) centered on the frequency of interest, and (ii) unevenly anchored.

#### Assessment

We visualized the simulation results as heat maps, showing the phase which was estimated to exhibit a maximal MEP after filtering.

## Results

### Simulation study 1

Our first simulation contrasted a narrow-band centered filter of high order followed by Hilbert transformation (approach 1) with a broad-band unevenly anchored filter of lower order followed by Fourier transformation (approach 2). It suggested that the second approach achieves better recovery of the original dependency. The first approach was unable to recover a unimodal phase-dependency of MEP amplitudes, while the second was able to discern between uni- and bimodal phase dependencies (see figure 1 B/C). Only when the order of the narrow-band filter was decreased did this approach recover a unimodal phase dependency (see figure 1 D) and distinguish it from a bimodal phase dependency (see figure 1 E).

### Simulation study 2

This simulation explored how the estimation of the optimal phase in the case of unimodal dependency was affected by various parameters of the signal processing pipeline. It highlighted that the phase, whether estimated with Hilbert (see figure 2 A/C) or Fourier transformation (see figure 2 B/D), appears to exhibit similar profiles. The main issue comes with higher filter orders by starting to distort the signal; a phenomenon which appears to start earlier for more narrow bands. At the same time, running a causal filter with an uneven bandwidth can bias the optimal phase estimate significantly (see figure 2 C/D).

**Figure 2.**
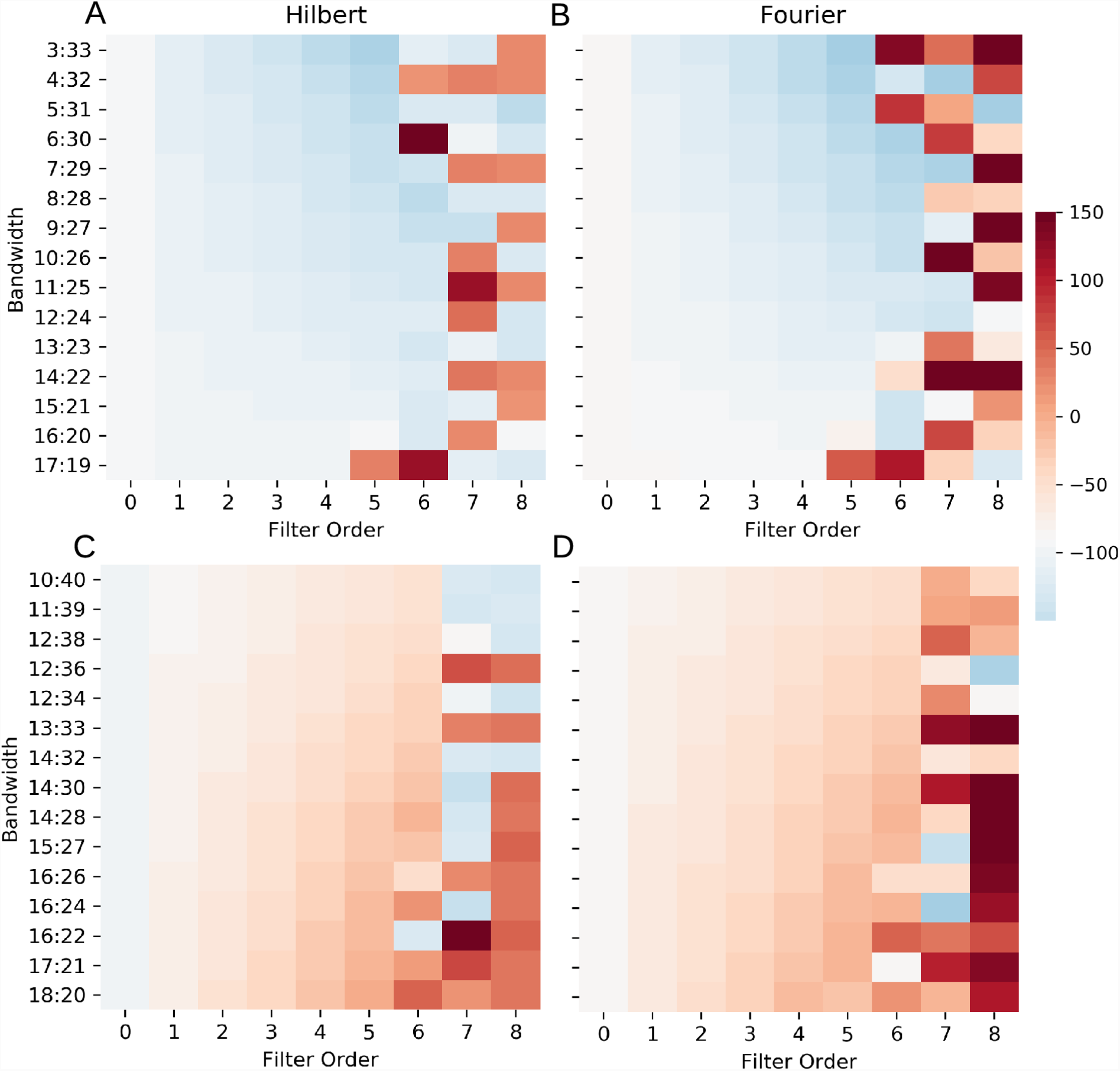
Estimated optimal phase for the unimodal dependency depends on filter width, center and order. All heat maps show, with filter order on the x-axis and bandwidth on the y-axis, the estimated optimal phase color-coded with a divergent colormap. The colormap is anchored with white to −90, i.e., the optimal phase according to the simulation. The rows show the recovery depending on whether Hilbert (A/C) or Fourier (B/D) transformation was used for phase estimation. The columns show the recovery for evenly centered (A/B) or uneven bands (C/D).

## Discussion

We conducted two simulation studies to explore how differences in signal processing affect the estimation of phase-dependency. The first of these studies applied two approaches based on earlier research (Keil et al., 2013; van Elswijk et al., 2010). The second explored the influence of processing on the phase estimation. The goal was to investigate how differences of data processing can explain supposedly contradictory findings in earlier literature, and explore the related pitfalls.

### Methodological considerations

One key result of our simulation studies was that a low filter order is required to recover uni- and bimodal phase-dependency patterns. If filter orders are too high, the signal becomes distorted, which in turn leads to an artifactual bimodal dependency (see figure 1 D/E). Given a sufficiently low order, both broad-band Fourier and narrow-band Hilbert transformation were able to recover the original phase-dependent relationship. The apparently bimodal dependency introduced by filtering is mainly artifactual, given that too high filter orders can corrupt the phase spectrum of a signal (Oppenheim et al., 2014).

At the same time, applying causal filters with uneven pass-bands induces phase-shifts. An evenly centered passband should therefore be considered as a step towards minimizing phase distortion, should a specific frequency be of interest.

More generally, if Hilbert transformation is used for phase-estimation, this can clash with the goal of estimating the phase for a narrow frequency band of interest. In such cases, phase estimation using Fourier transformation after broad-band low-order filtering might constitute a more suitable approach.

It should also be noted that using a temporal lag in relationship to the MEP necessarily introduces a phase-shift (see figure 1 D/E). For example, 5 ms at a frequency of 18 Hz correspond to ~32°. Correcting for this can be challenging. If the frequency is determined a priori, correcting the bias may be a good alternative. However, if more non-stationarity is required, unsupervised phase prediction, e.g., the use of an adaptive autoregressive model (Zrenner et al., 2017) might also be considered as a feasible option.

### Toolbox selection

Implementations of the same processing pipeline in different environments, e.g., by using different toolboxes or programming languages, can influence the results (Widmann et al., 2015). Notably, recent versions of a commonly used data analysis software (Fieldtrip, Oostenveld et al., 2011) support an automatic filter instability correction and interrupt the computation (e.g., “*fail with an exception*”), if the filter order is too high. This behavior was introduced in early 2013 (according to git log–grep “instability”, commit #914d6ab, https://github.com/fieldtrip/fieldtrip/); analyses performed prior to this software update might therefore not have experienced any error warning if the applied filter orders were too high.

## Conclusions

Findings can become biased by too high filter orders or uneven pass-bands. On a more general account, these observations propose two approaches: (i) Using up-to-date toolboxes and re-running scripts after software updates (Kitzes et al., 2018) and (ii) Running simulation studies in parallel to the actual data processing to safeguard the analysis pipeline against potential pitfalls (Haufe, 2015).

## Acknowledgments

M.S. was supported by the Graduate Training Centre of Neuroscience & International Max Planck Research School, Graduate School of Neural Information Processing, Tuebingen, Germany. A.G. was supported by grants from the German Federal Ministry of Education and Research [BMBF 13GW0119B, IMONAS; 13GW0214B, INSPIRATION; 13GW0270B, INAUDITAS] and the Baden-Wuerttemberg Foundation [NEU005, NemoPlast]. The authors declare no competing financial interests.

